# Causal inference shapes crossmodal postdiction in multisensory integration

**DOI:** 10.1101/2025.07.02.662778

**Authors:** G Günaydın, JK Moran, T Rohe, D Senkowski

## Abstract

In our environment, stimuli from different sensory modalities are processed within a temporal window of multisensory integration that spans several hundred milliseconds. During this window, stimulus processing is influenced not only by preceding and current information, but also by input that follows the stimulus. The computational mechanisms underlying crossmodal backward processing, which we refer to as crossmodal postdiction, are not well understood. We examined crossmodal postdiction in the audiovisual (AV) rabbit illusion, in which postdiction occurs when flash-beep pairs are presented shortly before and shortly after a single flash or a single beep. We collected behavioral data from 32 participants and fitted four competing models: a Bayesian causal inference (BCI), a forced-fusion, a forced-segregation, and a non-postdictive BCI model. The BCI model fit the data well and outperformed all other models. Building on findings demonstrating causal inference during non-postdictive multisensory integration, our study shows that the BCI framework is also effective in explaining crossmodal postdiction. Observers accumulate causal evidence that influences their perception of preceding stimuli following a causal decision. Our study shows that crossmodal postdiction in the AV rabbit illusion is formed within a temporal window of multisensory integration that encompasses past, present, and future input, which can be effectively explained by the BCI framework.

## 2. INTRODUCTION

Imagine that you are walking down the street when you see a friend waving and yelling to get your attention from a distance. As you notice your friend, you may realize that you had been hearing their calls for some time but were not aware of them. In addition to the current calls, you then may become aware of the previously neglected calls. This example suggests that the processing of current information can retroactively influence the perception of past stimuli that are still available in sensory memories, an observation that has often been described as postdiction (1,2). So far, the computations underlying postdiction have primarily been investigated in unisensory processing (3). In everyday life, however, stimuli typically originate from different sensory modalities, and they must be integrated or segregated to achieve a coherent perception of the multisensory environment (4,5). In a multisensory context, a major challenge for the brain is to determine whether to integrate or segregate information from different sensory modalities. Specifically, the brain must infer causality between the different sensory channels so that sensory information arising from a common cause can be integrated across modalities, with the weights proportional to the relative sensory precision, as expressed in the Bayesian causal inference (BCI) framework (6–8) (Fig. 1, top panel). Similarly, different sensory modality stimuli should be segregated if they have independent causes. In this causal inference framework, internal prior expectations are combined with the likelihoods - the incoming sensory evidence from the external stimuli - to infer causality and perceive latent properties of the environment according to their causal structure. While the BCI model accounts well for non-postdictive multisensory integration and segregation (15), it is currently unknown whether the BCI model can explain the computational mechanisms underlying crossmodal postdiction.

**Figure 1.**
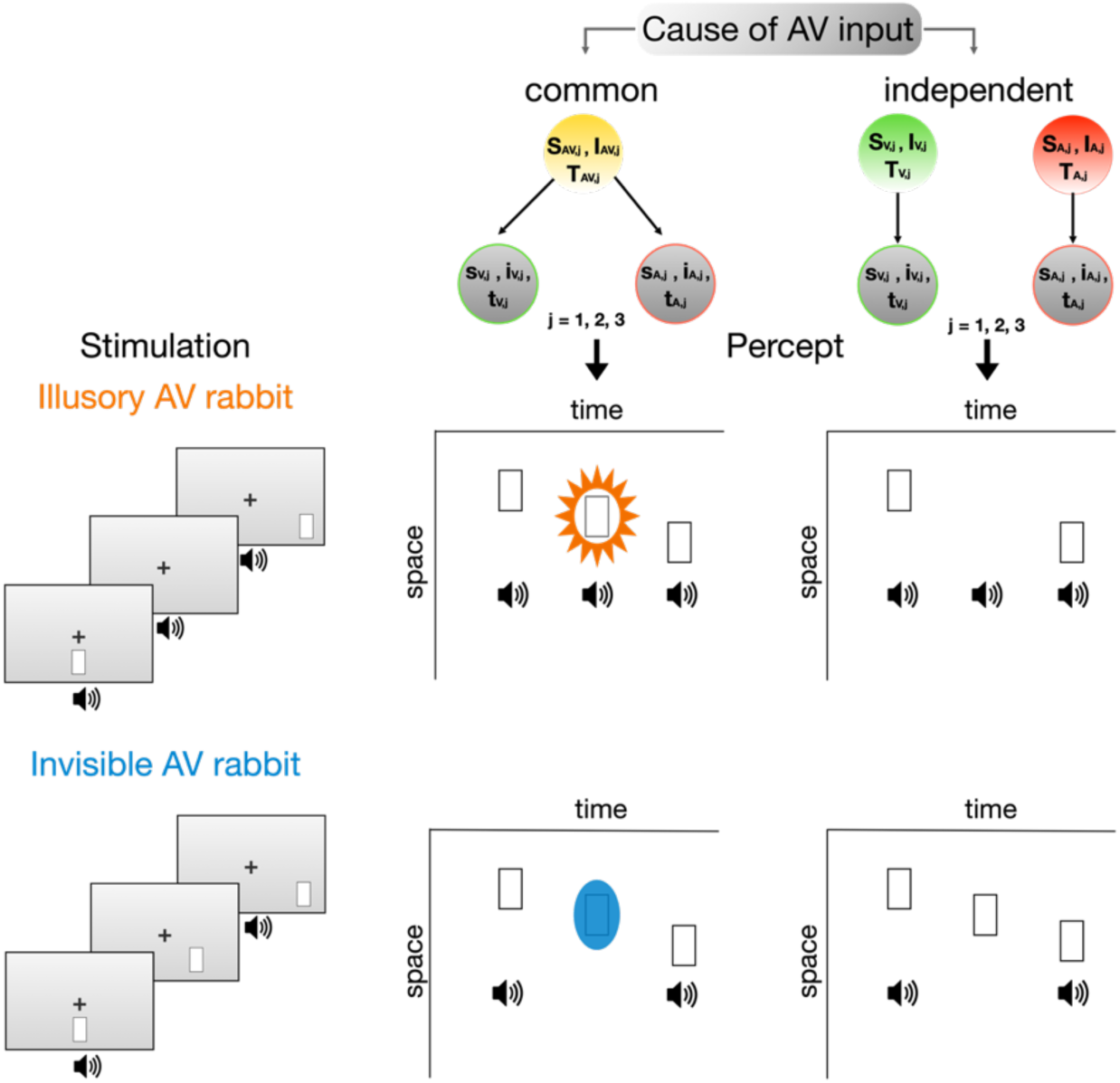
Audiovisual stimuli sequences in the illusory and invisible AV rabbit and perceptual outcome predictions of the generative BCI model. The figure illustrates an experimental trial in which the flash sequence moved from the center to the right. Participants were instructed to ignore the auditory stimuli and to report each perceived flash and its location. The Bayesian framework (top) posits that observers infer whether the auditory and visual inputs originated from two potential causal structures: common cause or independent causes. Observers combine causal priors with accumulating sensory evidence across time (t), space (s) and intensity (i) cues for the three different stimulus pairs (A, V, or AV, j = 1,2,3) to infer causality. If observers infer a common cause, they integrate the AV cues, leading to AV rabbit illusions. In the *illusory AV rabbit* (left, middle), a flash-beep pair is followed by a single beep, and then a second flash-beep pair. When observers infer a common cause of the AV inputs, this sequence of stimuli can result in an illusory flash that is localized in between the first and the second flashes (middle of middle panel, highlighted in orange). This indicates postdictive crossmodal processing, wherein the last flash-beep pair retroactively influences the perceived location of the illusory flash. In the *invisible AV rabbit* (left, bottom) a flash-beep pair is followed by a single flash, and then a second flash-beep pair. If observers infer a common cause of the AV inputs, this sequence of stimuli can result in an invisible flash, i.e., a flash that is perceptually suppressed (middle of bottom panel, highlighted in blue). Both illusions indicate crossmodal postdiction.

Previous studies using the BCI framework have improved our understanding of the computations underlying multisensory integration ((9–11). For example, the sound-induced flash illusion, wherein a single flash that is presented alongside two rapid tones, is perceived as two flashes, has been found to involve causal inference (12,13). A modified version of this illusion includes postdictive crossmodal effects: the *audiovisual (AV) rabbit illusion* (14). In the AV rabbit illusion, flash-beep pairs are presented shortly before and shortly after a single beep or a single flash. When observers perceive the AV rabbit illusion, an illusory single flash appears to be spatially localized between the first and the last flashes (Figure 1, *middle panel*). In contrast, in the so-called *invisible AV rabbit illusion*, a flash is perceptually suppressed (Figure 1, *bottom panel*). The BCI framework (7,15) offers a promising approach to account for postdictive crossmodal perception (16–18). To determine whether flash-beep sequences in the AV rabbit illusion arise from a common cause or independent causes, observers integrate causal evidence for each causal structure from all flash-beeps over time, space and signal presence. Flash-beep stimuli that fall within the temporal window of multisensory integration (17,19) increase the probability of a common cause. Specifically, the last flash-beep pair in the AV rabbit illusion provides additional sensory evidence to inform the causal probabilities. After accumulation of causal evidence, the brain subsequently integrates (or segregates) spatiotemporal information of the preceding stimuli in the case of a common cause, the flash-beeps are integrated with a stronger weight on the more salient auditory beeps (20). For the middle stimulus, integration of the single beep without a flash, leads to the perception of an illusory flash in the illusory AV rabbit because the brain infers an additional middle flash-beep event. In the invisible AV rabbit, integration of a single flash without a beep leads to the perceived absence of the flash because the brain infers that the middle flash-beep event did not occur. Because causal inferences are informed also by causal evidence from the last AV event, postdictive effects arise from the last flash-beep on the middle event. Hence, if the last flash-beep increases the probability of a common cause, it should also postdictively increase the probability that the middle event is integrated. Taken together, if causal inference could explain the AV rabbit illusion, we would expect that the BCI model captures postdictive crossmodal processes better than alternative non-causal models.

In this study, we presented 32 observers with an extensive set of temporally synchronous or asynchronous AV rabbit stimuli, as well as stimuli with changing directionality (*See Methods*). To examine whether causal inference accounts for crossmodal postdiction in the AV rabbit illusion paradigm, we developed and fitted a BCI model to estimate the causal structure underlying the flash-beep sequences, which included postdictive information from the last flash-beep event. Furthermore, we compared the BCI model with a non-causal forced-fusion (FF) model and forced-segregation (FS) models. The FF and FS models assume that the flash-beep stimuli are integrated and segregated in a mandatory fashion without accounting for the stimuli’s causal structure. To test how the last flash-beep pair postdictively influences causal inference as an explanation for the AV rabbit illusion, we also compared the BCI model with a non-postdictive BCI model (BCI-NP) that ignored causal information from the last flash-beep event. This was a critical comparison to test how the BCI model implicitly explains crossmodal postdiction.

## 3. RESULTS

To investigate the computational mechanisms underlying the illusory and invisible AV rabbit we presented participants with flash-beep sequences and asked them to report each perceived flash and its location. Compared to the original setup (14), we added experimental conditions (See *Methods*) to enable model fitting and to further improve the validity of the paradigm. To reduce a behavioral response bias that might occur if the flash sequence always moved in one direction, e.g. from left to right, we added conditions in which the flash sequence changed its direction (e.g., the first flash was presented in the center, the second flash to the right, and the third flash to the left). Due to the presence of direction changing conditions, the position of the last flash was not predictable from the first or second’s flash location. In the original study, the visual flash stimuli never changed their direction within a trial and hence, participants could predict the direction of motion of the flashes based on the location of the first flash. This may have influenced perception (21) and contributed to a response bias. Furthermore, conditions with a temporal asynchrony between visual and auditory inputs were added to test the range of the temporal binding window for the illusory and invisible AV rabbit. Finally, we included multisensory (audiovisual) and unisensory (visual only) control conditions to improve the parameter estimations in the computational models.

We quantified the strength of the illusions as illusion rates, that is the proportion of trials in which a middle flash was perceived (Illusory Rabbit) even though it was absent, or a veridical middle flash was not perceived (Invisible Rabbit). Further, responses were only counted as illusions if the participants located the flashes on specific locations and in a certain order. For instance, for the illusory rabbit, a trial response was considered as an illusion only if the participant correctly located and reported flashes in the first and third position, as well as an extra illusory flash in the middle in the corresponding respective order (See *Methods*).

### Sequence direction does not influence illusion rates

To test whether the direction of the flash sequence influenced illusion rates, following normality checks (*See Methods*), Wilcoxon signed-rank tests or paired t-tests were implemented to compare illusion rates between the right and left motion conditions. Following Bonferroni correction (corrected p-value = 0.005, for 10 pairwise comparisons), there were no significant differences between left and right motion directions. Therefore, we combined left and right motion trials in the following analysis steps.

### Illusion rates are higher in illusion conditions compared to control conditions

In line with the original study (14), we compared the illusion rates for the two illusion conditions with unisensory and multisensory control conditions (Figure 2). Both illusion conditions showed significantly higher illusion rates compared to the unisensory and multisensory control conditions (Illusory Rabbit: against unisensory control, p = 3.75×10^-6^, z = 4.62, against multisensory control, p = 3.76×10^-6^, z = 4.62; Invisible Rabbit: against unisensory control, p = 1.03×10^-5^, z = 4.41; against multisensory control, p = 3.76×10^-6^, z = 4.62). Furthermore, the illusion rates were significantly higher for the unisensory compared to the multisensory control conditions for both illusions (Illusory Rabbit: p = 7.49×10^-5^, z = 3.96; Invisible Rabbit: p = 1.40×10^-4^, z = 3.81). These findings largely replicate the results of Stiles *et al*. (14).

**Figure 2.**
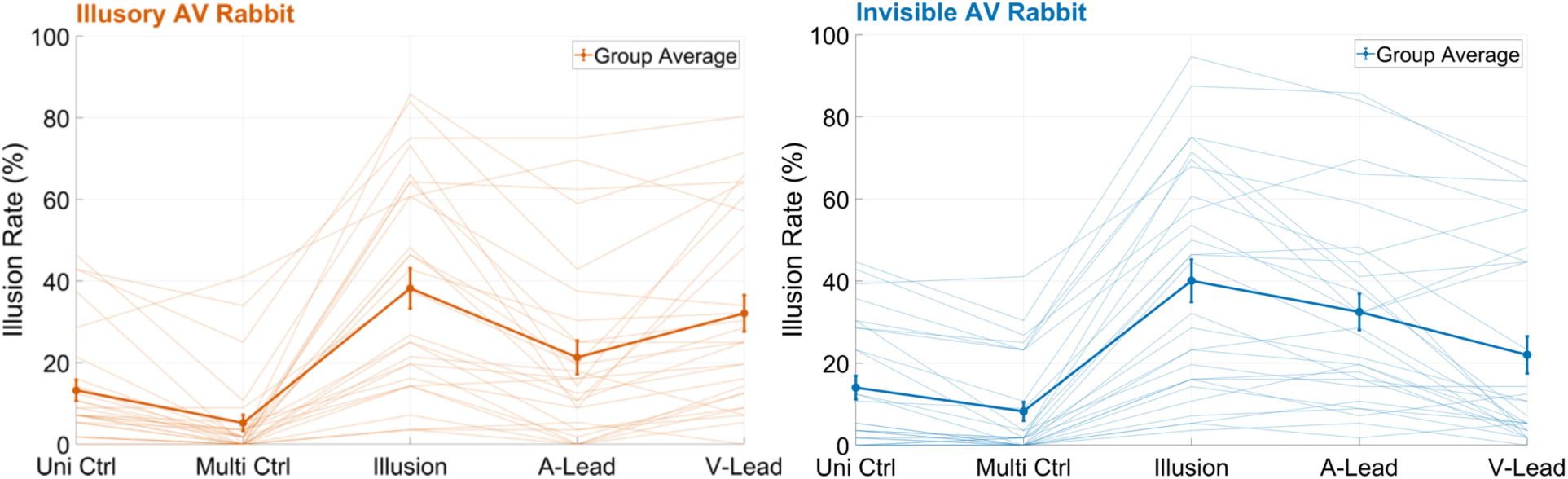
Illusion rates for the illusory and invisible AV rabbit are higher in the synchronous conditions than in the control and asynchronous conditions. The figure shows illusion rates for the unisensory and multisensory control conditions for the invisible (lower, blue) and the illusory (upper, orange) AV rabbit. Thicker opaque lines show the average across subjects and thinner translucent lines show individual data. The average illusion rates for the synchronous illusion conditions were around 40%, which were higher than the illusion rates in the multisensory and unisensory control conditions. Uni Ctrl = unisensory control; Multi Ctrl = multisensory control; A-Lead = auditory lead asynchronous condition; V-Lead = visual lead asynchronous condition.

### Audiovisual temporal asynchrony is associated with lower illusion rates

Next, we examined the effect of stimulus asynchrony between visual and auditory stimuli. We implemented two types of temporal asynchronies: (i) an auditory lead, in which the auditory stimulus sequence started 215 ms before the visual stimulus sequence, resulting in a 215 ms stimulus onset asynchrony the first auditory beep and the first visual flash (*Figure 4*), and (ii) a visual lead condition, in which the visual stimulus sequence started 215 ms before the auditory stimulus sequence. Nonparametric tests comparing the synchronous and asynchronous conditions revealed higher illusion rates for the synchronous conditions for both for the illusory AV rabbit (against A-Lead Asynchrony, p = 3.63×10^-5^, z = 4.13, against V-Lead Asynchrony, p = 0.01, z = 2.51) and the invisible AV rabbit (against A-Lead Asynchrony, p = 4.2×10^-3^, z = 2.86, against V-Lead Asynchrony, p = 1.21×10^-5^, z = 4.38). Thus, temporal asynchrony reduces both illusions, which is probably the case because the sequences of auditory and visual stimuli fall outside the AV temporal integration window in asynchronous conditions.

### BCI model outperforms the forced-fusion, forced-segregation and non-postdictive models

We fitted four computational models (BCI, FF, FS and BCI-NP) to the data. The main difference between the BCI, FF and FS models was the causal prior. Only the BCI model among the BCI, FF and FS estimated the stimuli’s causal structure from a causal prior and sensory evidence. In contrast, the FF and FS models assumed either a common cause leading to forced integration (i.e. equivalent to a causal prior fixed to 1) or independent causes leading to forced segregation (i.e. a causal fixed to 0), respectively. Next, we investigated how the sensory evidence of the third flash-beep pair was used postdictively in the BCI model to calculate posterior causal probabilities. To accomplish this, we excluded the posterior causal probability of the third flash-beep pair from the combination of posterior probabilities across all flash-beep pairs. Using this extended non-postdictive BCI model, we computed predictions about the flash reports to determine if sensory evidence from the last beep-flash pair modulates the illusory rates. Interestingly, the non-postdictive BCI (BCI-NP) model underestimated the rates of the AV rabbit illusion (Figure 3).

**Figure 3.**
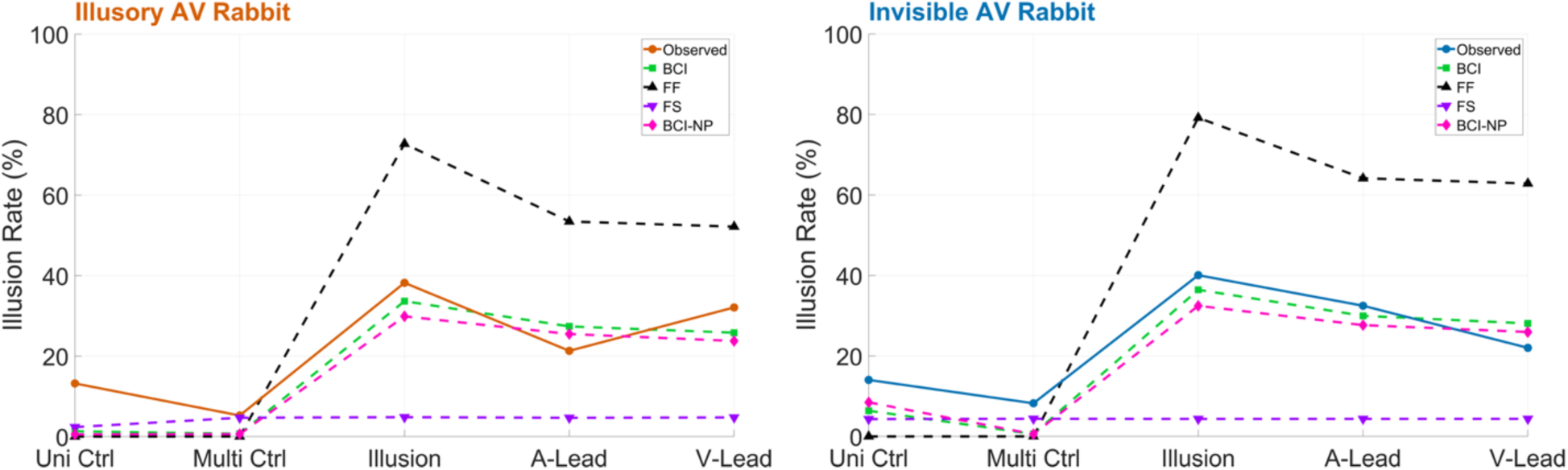
Model predictions demonstrate the BCI model’s superiority over other models in explaining the invisible and illusory AV rabbit. The group-averaged observed data (orange in the upper and blue in the lower panel) and the predicted illusion rates from the fitted BCI (green), forced-fusion (FF, black), forced-segregation (FS, purple), and non-postdictive BCI (BCI-NP, pink) models are shown for the invisible (lower) and illusory (upper) AV rabbit. Predicted illusion rates are shown for unisensory and multisensory control conditions, the illusion condition, and for asynchronous AV conditions. The BCI model shows the best overall data fit, as demonstrated by Bayesian model comparison (Table 1). Uni Ctrl = unisensory control; Multi Ctrl = multisensory control; A-Lead = auditory lead asynchronous condition; V-Lead = visual lead asynchronous condition

**Table 1.** Model comparison metrics for the BCI, FF, FS and BCI-NP models show strong superiority of the postdictive Bayesian causal inference (BCI) compared to a forced-fusion (FF), forced-segregation (FS), and non-postdictive BCI model. . Bayesian information criterion (BIC) and Akaike information criterion (AIC) quantify the trade-off between model fit (i.e., good explanation of the data) and model complexity (i.e., a sparse number of parameters). At group level, participant-specific relBIC_Group_ and relAIC_Group_ are summed over all participants. Lower values imply a better fit for a model. Nagelkerke’s coefficient of determination (R^2^) measured the proportion of explained variance against a null model of random guesses across 6 response options (i.e., across-participant mean reported). In a random-effects Bayesian model comparison, the protected exceedance probability (PEP) quantifies the probability that one model is more prevalent than competing models at group level, beyond which model frequencies could arise by random variations.

| MODELS | relBIC<sub>Group</sub> | relAIC<sub>Group</sub> | R<sup>2</sup> | PEP |
| --- | --- | --- | --- | --- |
| BCI | 0 | 0 | 0.94 | 0.96 |
| FF | 1938 | 1944 | 0.75 | 0 |
| FS | 572 | 577 | 0.88 | 0 |
| BCI-NP | 91 | 95 | 0.93 | 0.04 |

Model comparisons revealed that the BCI model outperformed the non-causal FF and FS models (Table 1), including the BCI-NP model, with respect to lower average Bayesian Information Criterion (BIC) and Akaike Information Criterion (AIC) values, as well as higher R^2^ values. Bayesian model selection revealed that the BCI model best explained the group data, with a protected exceedance probability of 0.96. FF, FS had negligible probabilities, and BCI-NP model had a probability of 0.04. Visual inspection of the BCI model’s predictions for the AV rabbit illusion showed that the model could accurately reproduce the invisible and the illusory AV rabbits and perception in the control conditions, as well as the effect of AV asynchrony (Figure 3). In contrast, the competing models overestimated (FF model) or underestimated (FS and BCI-NP models) the illusion rates.

## 4. DISCUSSION

In this study, we examined the computational mechanisms underlying crossmodal perception in the AV rabbit illusion. Our analysis revealed three main findings. First, we observed higher illusion rates in the two illusion conditions compared to unisensory and multisensory control conditions, even when experimentally controlling for potential response biases. Second, we found that temporal asynchrony between auditory and visual inputs diminished the illusion rates of both illusions. Third, we showed that the BCI model is more effective than other non-causal models at predicting the two illusions.

### Postdiction in the illusory and invisible AV rabbit

Our finding of higher illusion rates in the two illusion conditions compared to the unisensory and multisensory control conditions replicates the results of the original report by Stiles et al. (14). This suggests that the illusory and invisible AV rabbit illusions involve postdictive crossmodal processing. In the current study, we took great care to reduce a possible behavioral response bias that could have influenced the results of the original study. In that study, the visual flash sequence never changed its direction within a trial, which enabled top-down predictions of the location of subsequent flashes. In our study, the first flash was always located in the center of the screen. This prevented spatial predictions about the upcoming flash sequence. In addition, we included trials in which the flash sequence changed its direction with equal probability. These modifications should have significantly reduced the influence of sequence predictions and the possible resulting response bias. Therefore, we propose that a behavioral response bias is unlikely to account for the postdictive crossmodal effects observed in the current study.

### Temporal asynchrony reduces illusion rates in the illusory and invisible AV rabbit

To investigate how temporal asynchrony influences the two illusions, we presented conditions in which the visual and auditory stimulation sequences were asynchronous by 215 ms. Compared to synchronous trials, asynchronous trials resulted in fewer illusion reports for both illusions. Postdictive effects are stronger when the respective sensory inputs are closely aligned within the temporal window of multisensory integration. In other words, postdictive crossmodal effects are reduced when the inputs fall within the periphery of the window, which can last up to several hundred milliseconds (22,23). Preliminary data from Stiles et al. (18,24) suggest that the AV rabbit illusion can be found even when the final flash-beep pair is delayed by up to 500 ms; however, the illusion rates are substantially reduced in this case. The current results, in which asynchronies of 215 ms between the sensory stimulation sequences were used, support this finding. Interestingly, we observed greater inter-subject variability among subjects in the asynchronous conditions compared to the synchronous conditions. Furthermore, the effect of AV asynchrony was not symmetric for auditory versus visual lead, and this effect was different for the illusory compared to the invisible rabbit illusion (Figure 2). This might be related to the observation that asynchronies are handled differentially in different multisensory illusion paradigms (25). Taken together, our results and the preliminary data by Stiles et al. (18,24) demonstrate that a temporal window of multisensory integration exists, lasting a few hundred milliseconds, during which stimuli can retroactively influence the processing and perception of preceding stimuli, even if these stimuli occur in different sensory modalities.

### Bayesian causal inference effectively accounts for crossmodal postdiction

Consistent with studies on non-postdictive multisensory processing (10,11,26,27), we fitted three models to the data: FF, FS, and BCI. Across the experimental conditions, the BCI model outperformed the other two models. Our results align with those of studies on the sound-induced flash illusion (13), which has a design similar to that of the AV rabbit illusion but does not explicitly incorporate crossmodal postdiction. Shams et al. (12) showed that the sound-induced flash illusion can be adequately explained by a BCI framework that posits an observer who optimally combines auditory and visual inputs. Our results extend this finding by demonstrating that causal inference can account for crossmodal postdiction as well.

The Bayesian framework has previously been applied to elucidate the computations underlying postdiction in unisensory paradigms, such as the flash-lag effect (28) and the cutaneous rabbit illusion (3). In these paradigms, a unisensory low-speed prior has been found to play an important role, contributing to both prediction and postdiction. In the current study, we did not model the perception of the AV sequences using speed priors and likelihoods, but rather modeled independent AV events that are integrated if an observer infers a common cause for all events. Specifically, we propose that inferring a common cause of audiovisual stimuli can generate illusory percepts retrospectively in the illusory and invisible AV rabbit paradigm. The brain’s perception of the causal structure of AV events is determined by computing the spatiotemporal and numerical disparity of these events. Thus, the decision to integrate AV information occurs only after the entire AV stimulus sequence has been presented, paving the way for crossmodal postdiction. More specifically, causal inferences are informed by sensory causal evidence across all flash-beep pairs, including the last flash-beep pair. Thus, postdictive effects of the last flash-beep on the middle event will occur through causal inference: if the last flash-beep increases the likelihood of a common cause, it retroactively increases the likelihood that the observer will infer an AV middle event, leading to the AV rabbit illusions. This is because the beeps and flashes are integrated by weighing their relative sensory precision in case of a common cause inference. Due to the highly salient and precise auditory stimuli in the current paradigm, the auditory information can induce the illusory perception of a flash, which is also the case in the sound-induced flash illusion (13,29,30). The veridical number of flashes is perceived without postdictive influence when independent causes are inferred for the AV information, i.e., when it is processed separately (Figure 1, *left panel*).

Critically, our model accumulates sensory causal evidence across all AV events: Not only the last flash-beep pair but also the first AV event provides sensory causal evidence. Thus, the AV rabbit illusion also predictively dependents on the first flash-beep. In other words, prediction and postdiction emerged naturally from our BCI model. To disentangle postdictive from predictive effects, we fitted a BCI model in which we reduced the postdictive component of the causal inference process, i.e., by eliminating the sensory causal evidence from the last flash-beep pair. Interestingly, the non-postdictive model underestimated the illusion reports, and it was inferior to the full BCI model. This suggests that the illusion decisively involves crossmodal postdictive interactions via causal inferences related to the processing of the last flash-beep pair. However, the non-postdictive model captured at least parts of the AV rabbit illusions, suggesting that non-postdictive processes, such as prediction from the first AV event, also contribute to the illusions. In summary, our data show that the illusory and invisible AV rabbit illusions can be well described in terms of the BCI model. Similar to what has been previously shown for non-postdictive processing of multisensory stimuli (9–11,26,31) and for postdictive unisensory processing (3,32), our study provides evidence that crossmodal postdictive processing can be effectively accommodated by the BCI model.

### Neural dynamics and mechanisms that could underlie crossmodal postdiction

An open question concerns the precise neural mechanisms underlying causal inference and crossmodal postdictive processing. According to a hierarchical model of causal inference in the brain (10,11,33) and in line with predictive coding (34), prefrontal cortical areas may accumulate sensory causal evidence across the AV sequence via recurrent message passing across the cortical hierarchy to make a causal decision (11,26,33). Accordingly, posterior multisensory regions integrate or segregate the AV representations to compute AV estimates that account for the most likely causal structure. Thus, the postdictive effect of the last AV event may manifest as an accumulation process of sensory causal evidence. This would be followed by a final perceptual estimate that may include an illusory or invisible AV flash, if the probability of a common cause is high. Recently, Stiles et al. proposed a model of re-entry postdictive processing in the visual cortex for the AV rabbit illusion (18). In this model, the processing of the illusory flash, generated by the first flash-beep pair and the subsequent beep, is maintained long enough in the visual cortex to overlap with the processing of the second flash-beep pair. This reallocates the illusory flash between the first and the third flashes. It is also possible that such re-entry processing relies on multi-timescale oscillatory brain dynamics, including feed-forward and feedback processing between sensory and higher-order association cortices (5). Finally, ongoing neural oscillations, particularly in the alpha band (8-12 Hz), may influence the integrative processing of auditory and visual stimulus sequences. Prestimulus alpha oscillations have been associated with crossmodal integration (35–37) and postdiction in unisensory processing (38). Taken together, the results of the current study allow us to draw concrete assumptions about the neural dynamics underlying the illusory and invisible AV rabbit. Future studies could test these assumptions using functional neuroimaging or electrophysiological methods.

### Summary and conclusion

Previous studies demonstrated the effectiveness of the BCI framework in elucidating the computations underlying non-postdictive multisensory integration and postdiction in unisensory paradigms. Here, we applied computational modeling to the illusory AV rabbit paradigm, which combines these two areas of research, i.e. multisensory integration and postdiction. Our results show that the BCI framework effectively accounts for crossmodal postdiction and outperforms non-causal models and non-postdictive simulation extensions. Our study provides strong evidence that the BCI framework can be applied to the processing of stimulus sequences across the entire temporal window of multisensory integration, including crossmodal postdiction phenomena.

## 5. METHODS

### Participants

Thirty-two human volunteers (15 females, 16 males and one non-binary individual; mean age: 27y, ranging from 20y to 41y; one left-handed), participated in the study after providing written informed consent. Two participants were excluded from the analysis because they did not follow the instructions and could not complete the entire

experimental session. After the exclusion of these participants, two more participants were excluded as their illusion rates in the unisensory and multisensory control conditions were three standard deviations higher than the group average. Data from 28 participants (13 females, 14 males and one non-binary individual; mean age: 27y, ranging from 20y to 33y; one left-handed individual) were included in the final data analysis. Handedness was assessed using the Edinburgh Handedness Inventory. None of the participants reported a history of neurological or psychiatric disorders and all had normal or corrected-to-normal vision and hearing. The study was approved by the Ethical Committee of the Charité – Universitätsmedizin Berlin (approval number: EA2/039/19).

### Setup and stimuli

The experiment consisted of 3 training and 14 experimental blocks, which included visual-only and audiovisual stimuli. Prior to the experimental blocks, one additional block with auditory-only stimuli was presented to estimate the individual auditory precision. In this block participants had to report the number of perceived auditory beeps. Each of the 14 experimental blocks lasted around 5 minutes and between blocks, participants were allowed to take short breaks. At the beginning of the experiment, which had an overall runtime of about 2 hours, participants were provided with verbal and written instructions regarding the task. The study was conducted in a soundproof chamber, which was dimly lit. The presentation of auditory and visual stimuli was controlled using Psychtoolbox 3.09 (39). Visual stimuli (rectangle bars) were presented on VPixx’s VIEWPixx high quality 120 Hz calibrated research-grade screen. The flashes could appear at five locations, aligned horizontally at 4° below a central fixation cross. The locations were (from left to right): -5.68°, -2.84°, 0°, 2.84°, 5.68° visual angles. Flashes (of luminance 54.1 cd/m^2^) were presented against a gray background (of luminance 8.7 cd/m^2^) with a width of 0.56° visual angle and a height of 2.4° visual angle. Each flash was presented for about 17 ms. Auditory stimuli (beeps) were presented through a loudspeaker located centrally below the display. They consisted of a square wave tone with a carrier frequency of 800 Hz and a sound pressure level of 88 dB. Each beep lasted 7 ms.

### Experimental paradigm

The illusory and invisible AV rabbit paradigms were adapted from Stiles *et al*. (14). To allow for the modeling of postdictive crossmodal perception, we extended the original study by various stimulation conditions (Table 2). The participants’ task was to ignore auditory stimuli and to report each perceived flash and its location in each trial by successive button presses (after each trial) on a keyboard with five response keys. The response keys on the ergonomically distributed keyboard represented the five horizontal locations where a flash could appear on the screen. To ensure that participants kept their gaze on the central fixation cross, catch trials in which the cross turned into a circle (for 100 ms, 150 ms after the beginning of a trial) were presented in 6.66% of all trials. The group average performance for the fixation catch trials was 98.7% ± 0.56 (Standard error of the mean). Participants were instructed to press the middle key when this happened. Participants were also informed that most trials contain 2 or 3 flashes, that the first flash would always be presented at the central location, and that the sequence of flashes could move in one direction (left or right) or could change direction.

**Table 2.** Overview of visual and audiovisual experimental conditions. Stimuli are categorized for the 28 experimental conditions, from left to right, according to the number of flashes and beeps, the locations of the flashes on the screen, and the postdictive illusory reports taken into account for the behavioral analysis. For a trial report to be regarded postdictive, locations and order are essential, regardless of the number of flashes reported. For the conditions, R and L represent the right and left sequences, respectively. The “1-5” represent the locations on the screen with position 3 in the middle, 4 and 5 to the right and 1 and 2 to the left. A-Lead represents the auditory lead and V-Lead represents the visual-lead asynchrony conditions. Type 1 and type 2 are two different combinations of direction-changing conditions. 8 of the 18 conditions with a second flash presented the flash with a change of direction. Thus, the regularity of the sequence was only probabilistically predictable from the second flash. This reduced a potential explicit response bias for the illusory AV rabbit which could have occurred with only continuous center-to-periphery sequences: if a second (illusory) flash had been seen, it could have occurred on the left (2) or right (4) position. Twelve of the 28 conditions were asynchronous. In synchronous trials, the 7 ms auditory beeps and the 17 ms visual flashes for a given pair of flash-beeps were presented simultaneously and their stimulus onset was 75 ms apart from the following pair of flash-beeps. However, in two types of asynchronies, either auditory beeps or visual flashes were presented first, referred to here as auditory lead and visual lead, respectively (Figure 4). Thus, only three inputs of a single modality were presented first, with a stimulus onset of 75 ms between them, and after 58 ms the sequence of the other modality was presented.

| Condition | Flash-Beep Pair 1 |  | Flash-Beep Pair 2 |  | Flash-Beep Pair 3 |  | Veridical Flash Locations | “Illusory” Report Sequence |
| --- | --- | --- | --- | --- | --- | --- | --- | --- |
|  | Auditory | Visual | Auditory | Visual | Auditory | Visual |  |  |
| Invisible Rabbit Multisensory Ctrl, R | + | + | + | + | + | + | 3-4-5 | 3-5 |
| Invisible Rabbit Multisensory Ctrl, L | + | + | + | + | + | + | 3-2-1 | 3-1 |
| Invisible Rabbit Unisensory Ctrl, R | - | + | - | + | - | + | 3-4-5 | 3-5 |
| Invisible Rabbit Unisensory Ctrl, L | - | + | - | + | - | + | 3-2-1 | 3-1 |
| Invisible Rabbit, R | + | + | - | + | + | + | 3-4-5 | 3-5 |
| Invisible Rabbit, L | + | + | - | + | + | + | 3-2-1 | 3-1 |
| Illusory Rabbit Multisensory Ctrl, R | + | + | - | - | + | + | 3-5 | 3-4-5 |
| Illusory Rabbit Multisensory Ctrl, L | + | + | - | - | + | + | 3-1 | 3-2-1 |
| Illusory Rabbit Unisensory Ctrl, R | - | + | - | - | - | + | 3-5 | 3-4-5 |
| Illusory Rabbit Unisensory Ctrl, L | - | + | - | - | - | + | 3-1 | 3-2-1 |
| Illusory Rabbit, R | + | + | + | - | + | + | 3-5 | 3-4-5 |
| Illusory Rabbit, L | + | + | + | - | + | + | 3-1 | 3-2-1 |
| Illusory Rabbit, A-Lead, R | + | + | + | - | + | + | 3-5 | 3-4-5 |
| Illusory Rabbit, V-Lead, R | + | + | + | - | + | + | 3-5 | 3-4-5 |
| Illusory Rabbit, A-Lead, L | + | + | + | - | + | + | 3-1 | 3-2-1 |
| Illusory Rabbit, V-Lead, L | + | + | + | - | + | + | 3-1 | 3-2-1 |
| Invisible Rabbit, A-Lead, R | + | + | - | + | + | + | 3-4-5 | 3-5 |
| Invisible Rabbit, V-Lead, R | + | + | - | + | + | + | 3-4-5 | 3-5 |
| Invisible Rabbit, A-Lead, L | + | + | - | + | + | + | 3-2-1 | 3-1 |
| Invisible Rabbit, V-Lead, L | + | + | - | + | + | + | 3-2-1 | 3-1 |
| Direction Changing, 1 | + | + | + | + | + | + | 3-4-1 | Not defined |
| Direction Changing, 2 | + | + | + | + | + | + | 3-2-5 | Not defined |
| Direction Changing Unisensory, 1 | - | + | - | + | - | + | 3-4-1 | Not defined |
| Direction Changing Unisensory, 2 | - | + | - | + | - | + | 3-2-5 | Not defined |
| Direction Changing, A-Lead, 1 | + | + | + | + | + | + | 3-4-1 | Not defined |
| Direction Changing, V-Lead 1 | + | + | + | + | + | + | 3-4-1 | Not defined |
| Direction Changing, A-Lead, 2 | + | + | + | + | + | + | 3-2-5 | Not defined |
| Direction Changing, V-Lead, 2 | + | + | + | + | + | + | 3-2-5 | Not defined |

Pilot data have shown that these instructions reduce the response bias that some subjects showed without these instructions. In the experimental runs stimuli consisted of sequences of 2 or 3 flashes, which were presented with stimulus onset asynchronies of 75 ms (Figure 4). Flashes were presented together with 0, 2 or 3 beeps that were not task relevant. In total, there were 28 experimental conditions (Table 2) and each participant completed 28 trials per condition. In each trial participants had a response window of 3 seconds. The inter-stimulus interval between trials varied randomly between 1.2 and 1.8 seconds (mean = 1.5 seconds). Hence, each trial lasted 4 to 5 seconds. In each block, participants completed 60 trials, including four catch trials and the order of the 28 experimental conditions was randomized across blocks.

**Figure 4.**
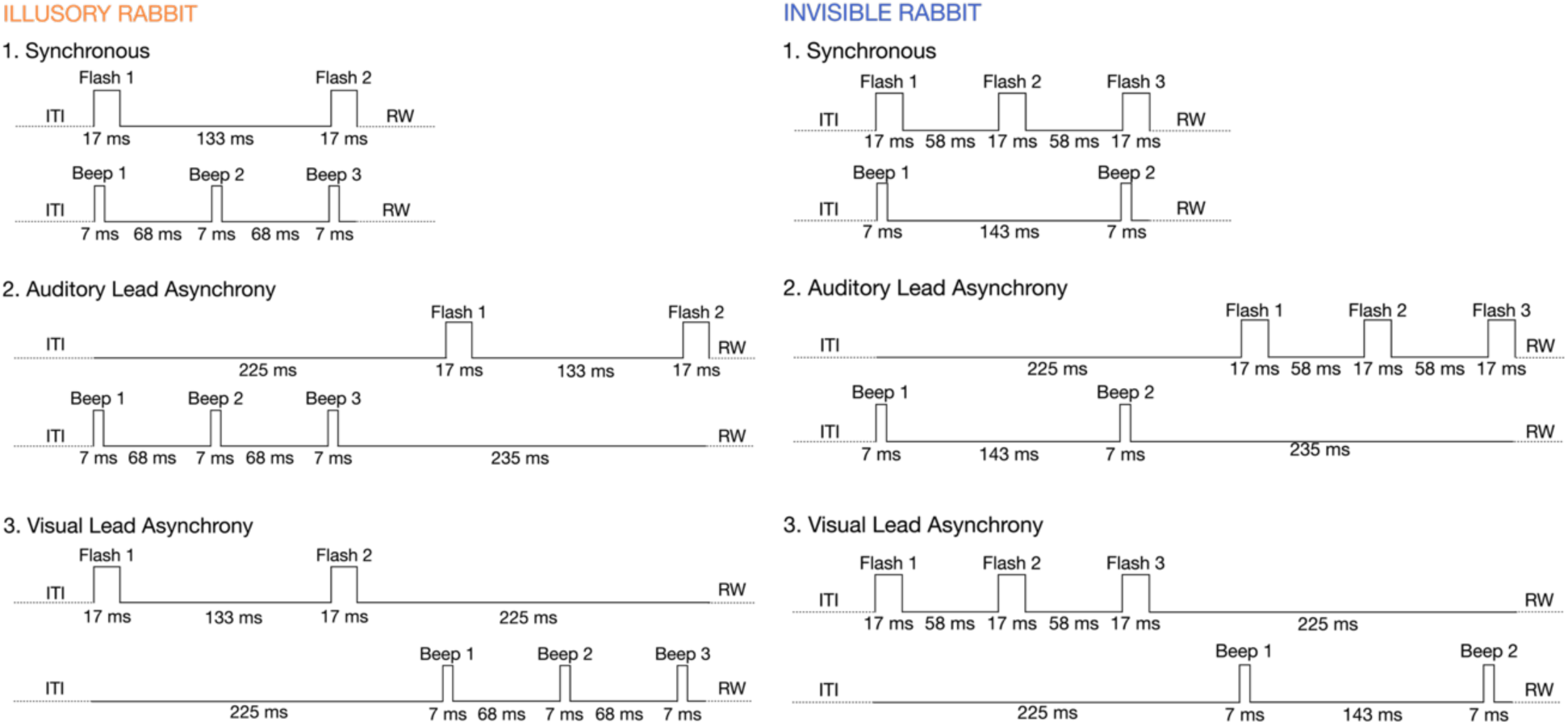
Timing of auditory and visual inputs in illusory and invisible AV rabbit trials. The figure illustrates the input timing for the synchronous AV trials (top), auditory lead AV trials (middle), visual lead trials (bottom) for the invisible AV rabbit (upper) and the illusory AV rabbit (lower). ITI = intertrial interval; RW = response window.

### Data analysis

Data analysis was performed, and plots were generated using MATLAB version R2024a (40). Participants’ flash reports were analyzed by computing AV rabbit illusion rates. A trial response was regarded as an illusion for the illusory AV rabbit if participants reported three consecutive flashes in the first, second and third locations (i.e., 3-4-5 or 3-2-1). For the invisible AV rabbit, a trial response was considered an illusion if participants reported two consecutive flashes in the first and third locations (i.e., 3-5 or 3-1). Illusory reports for each experimental condition are defined in Table 2. In the control trials, a “match/hit” was defined as correctly reporting the veridical order of the visual flashes on the screen. For each participant and condition, the percentage of correct responses based on illusory reports or match responses (reporting the veridical sequence) was calculated. First, Shapiro-Wilk tests were performed to investigate the normality of the data distribution. Because most of the illusion rates violated the normality assumption, the effects flash sequence direction on the illusion rates was tested with nonparametric (Wilcoxon signed rank) tests. There was no evidence for an effect of direction (left to right, or right to left). Since direction effects were not part of our research question, data from left and right directions were combined. This resulted in ten experimental conditions for the further data analysis. For each illusion condition, the main research question was whether the illusion rates differed between the illusory conditions and the unisensory and multisensory control conditions, as well as the asynchronous illusion conditions. To this end, nonparametric tests Wilcoxon signed-rank tests were computed. All p-values were adjusted using Bonferroni correction for multiple comparisons.

### Computational modelling of the behavioural data

To investigate whether participants adopted a causal inference or an alternative non-optimal decision strategy such as forced fusion or forced segregation, four models (BCI, non-postdictive BCI, forced-fusion and forced-segregation) were fitted to the reports of flashes and their locations. Bayesian model comparison was used to determine which model best explained participants’ behavioral data and the predictions of the causal inference and competing models for the illusion rates were examined. The BCI model combined the precision-weighted fusion estimate with the auditory segregation estimate proportional to the posterior probability of a common or independent causes, respectively. The forced fusion model (6,16) integrated the AV stimuli, weighted by their relative precision in a mandatory fashion. Finally, the forced segregation model assumed that participants’ flash percepts were independent from any auditory inputs, i.e. beeps. Details on the BCI model and the fitting to audiovisual perception on singular dimension such as spatial location and stimulus number can be found in Koerding *et al*. (7) and Rohe *et al*. (7,10). For perception of multivariate cues such as our flash-beep sequences, the approach implemented by Samad et al. (41) was extended to signals described by location, timing and intensity cues (Figure 1). The absence of a signal was conceived as a stimulus of zero intensity.

Our generative BCI model assumed that common (C=1) or independent (C=2) causes were determined by sampling from a binomial distribution with the causal prior p(C=1) = p_common_. For a common cause, each of the three independent flash-beep pairs S_AV,j_ was drawn from a common trivariate normal distribution N(μ_P_, Σ_P_), which modeled a cue for location S_AV,j_ (along the azimuth), time T_AV,j_ (relative to fixation), and intensity I_AV,j_ (which can be above or below a perceptual threshold) of the flash-beep pair. We assumed independence of the three stimulus components, so that the trivariate normal distribution was characterized by the prior spatial, temporal, and intensity means μ_S,P, j,_ μ_T,P,j_, μ_I,P,j_ (separately for beeps (μ_I,P,A,j_) and flashes (μ_I,P,V,j_)) and the diagonal covariance matrix of the prior spatial, temporal, and intensity standard deviations σ_S,P_, σ_T, P_ and σ_I,P_ (i.e., the same for j ∈ {1, 2, 3}).

For two independent causes, the “true” distal auditory and visual locations (S_A,j_ and S_V,j_), timing (T_A,j_ and T_V,j_), and intensity (I_A,j_ and I_V,j_) of beeps and flashes were drawn independently from the prior distribution. The observer does not have direct access to the distal beep-flash sequences but needs to infer and estimate their properties from proximal noisy sensory inputs x_A,j_ = (s_A,j_, t_A,j_, i_A,j_) and x_V,j_ = (s_V,j_, t_V,j_, i_V,j_). Sensory noise was introduced by sampling the sensory inputs independently from trivariate normal distributions. These distributions were centered on the experimentally defined auditory (respectively visual) temporal and spatial location as well as intensity of the stimuli. If a stimulus was presented or absent in a sequence, the intensity was set to 1 or 0, respectively. We assumed independence of the sensory inputs’ cues so that the variance is characterized by a diagonal covariance matrix with parameters σ_s_A, σ_t_A, σ_i_A (respectively σ_s_v, σ_t_V, σ_i_V). Further, we assumed that an observer only reported a flash if its intensity iV,j was above a perceptual threshold Φ_I_. Overall, the basic generative model included the following 11 parameters: the causal prior p_common_, the prior’s spatial, temporal and intensity standard deviations σ_S,P_, σ_T,P_ and σ_I,P_, the auditory standard deviations σ_s_A, σ_t_A, σ_i_A, and the visual standard deviations σ_s,V_, σ_t,V_, σ_i,V_, and the perceptual intensity threshold Φ_I_. Please note that we denote all variables and parameters related to the distal stimuli with uppercase letters and all variables and parameters related to the proximal sensory inputs with lowercase letters. The prior’s spatial mean μ_S,P,j_ was fixed to zero for all three flash-beep pairs j, assuming that observers expect unbiased, centered audiovisual stimuli. The prior’s temporal mean μ_T,P,j_, were fixed to 47, 105 and 163 ms for j ∈ {1, 2, 3} as experimentally defined (Figure 4). Only for the intensity prior, we differentiated between a visual and auditory intensity prior to reflect different a priori likelihoods of stimulus occurrence which can be quickly learned by the observer across all experiment conditions: The prior’s intensity means μ_I,P,V,j_ was fixed to 1 for j ∈ {1, 3} and to 0.65 for j ∈ {2}, for the visual stimuli, whereas μ_I,P,A,j_ was fixed to 0.78 for j ∈ {1, 3} and to 0.5 for j ∈ {2}, for the auditory stimuli. These μ_I,P,V,j_ and μ_I,P,A,j_ values represent the proportion of trials across all conditions in which a stimulus was indeed presented in the experiment.

Because the observer does not have direct access to the generative process, the observer needs to infer the causal structure and perceptually estimate properties of the flash-beeps from sensory inputs: For the causal inference according to the full postdictive BCI model, we assume that the observer accumulates the causal sensory evidence across the three flash-beep events and combines the evidence with the causal prior: Given flash-beeps’ vectorial sensory inputs x_A,j_ and x_V,j_, the observer infers the posterior probability of the underlying causal structure by combining the causal prior with the sensory likelihood according to Bayes rule:

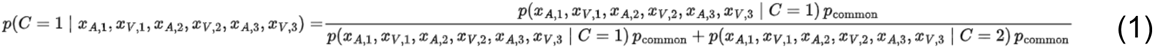

The sensory likelihood under a common cause assumption (C = 1) is accumulated over three independent flash-beeps and their independent spatial, temporal and intensity cues:

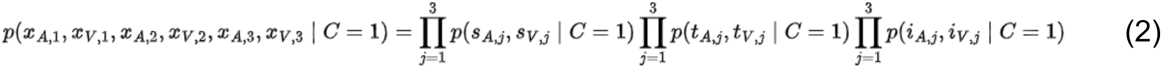

The sensory likelihood for each cue can be obtained by integrating over flash-beep pairs (7), e.g. for the spatial component S_AV,j_ (and likewise for the temporal and intensity components):

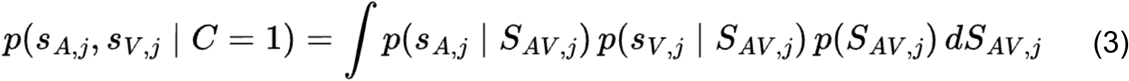

The three factors in the integral are Gaussian and can therefore be solved analytically:

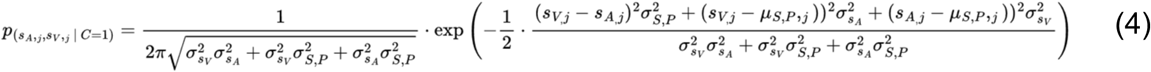

In an analogous fashion to equation 3, we can derive and compute the sensory likelihood under an independent cause assumption (C = 2):

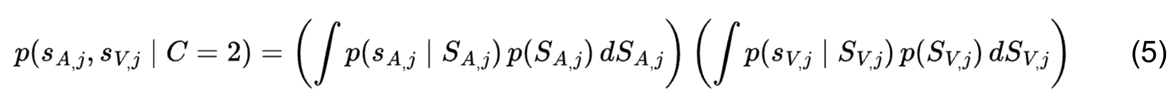

The Gaussian distributions allow for an analytic solution:

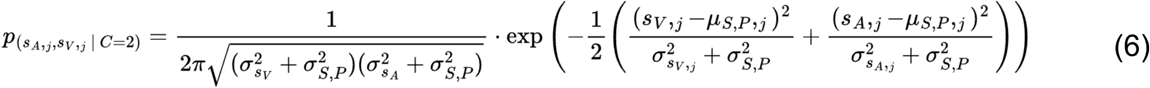

Effectively, the posterior probability of the underlying causal structure (Equation 1) combines the sensory likelihoods (Equations 4 and 6) with the causal prior: Critically, the sensory likelihoods under both causal structures are accumulated (i.e., mathematically multiplied) across the cues of all three flash-beep pairs according to equation (2).

For optimal perceptual inference of flash-beep estimates in the case of a common cause (C=1), the audiovisual estimate of a spatial, temporal or intensity of the stimulus (e.g., the spatial component S^$^_AV,C=1,j_) is obtained by combining the auditory and visual cues as well as the prior weighted by their relative reliability as quantified by the inverse of sensory variances (i.e., fusion estimate):

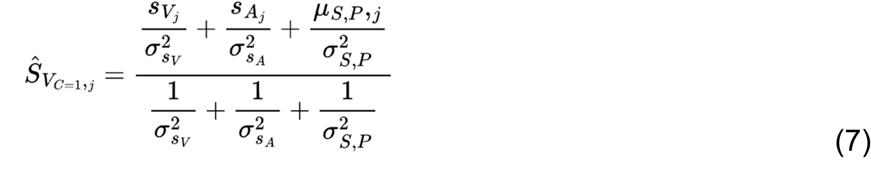

In the case of independent causes (C=2), the optimal estimates of the unisensory visual stimulus component (e.g., the spatial component S^$^_V,C=2, j_) are independent from the auditory cues (i.e. segregation estimate):

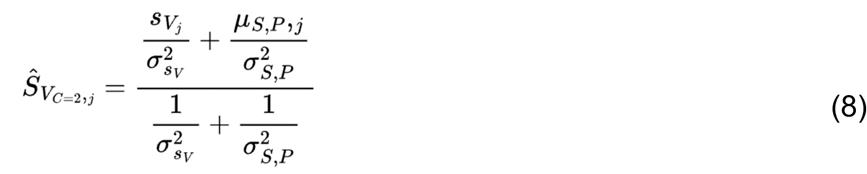

Because the observer needs to infer whether the sensory cues come from common or independent sources, the observer’s causal inference is subject to a uncertainty which also influences the observer’s perceptual inference: To account for observer’s causal uncertainty, the model computes final task-relevant spatial, temporal and intensity estimates of the distal stimuli (e.g. spatial S^$^_V_) by combining the fusion estimate (e.g., S^$^_AV,C=1,j_ for C=1) and the task-relevant segregation estimate (e.g., S^$^_A,C=2,j_ for C=2) depending on the posterior probabilities of the estimates’ underlying causal structures. In the ‘model averaging’ strategy of the BCI model, the observer weighs the estimates in proportion to the posterior probabilities of their underlying causal structures, for example for the posterior visual spatial estimate S^$^_V, j_:

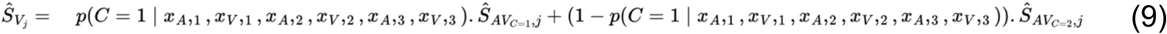

Note that the observer can also combine the numerical estimates according to different decision strategies such as model selection and probability matching (42). We compared the Bayesian ‘model averaging’ decision strategy, which take causal uncertainty into account, with two additional non-optimal heuristic strategies: First, a classical forced-fusion model (6) that mandatorily assumes a common cause (i.e., p_common_ = 1), thus always selecting the fusion estimate (Equation 7). Second, a forced segregation model that mandatorily assumes independent causes (i.e., p_common_ = 0), thus always selecting the segregation estimate (Equation 8). The forced fusion and segregation models both have one free parameter less (i.e., p_common_), thus reducing the complexity of model compared to the full BCI model.

Finally, we investigated a potential postdictive mechanism from the last beep-flash pair on the perception of the illusory and invisible rabbit by testing the influence of the sensory evidence of the third flash-beep pair on causal inferences in the BCI model. In the non-postdictive BCI model, we assumed that the observer accumulates the causal evidence only across the first and second flash-beep events but ignores the third event. Thus, we excluded the sensory evidence of the third flash-beep pair from the combination of sensory likelihoods across all flash-beep pairs (i.e., Equation 2, j ∈ {1, 2}).

To compare the four candidate models, we fitted each model to participants’ behavioral reports of the perceived number and location of flashes based on the predicted distributions of the visual estimates (i.e. the marginal distributions, e.g. for the location component: p(S^$^_A,j_ |S_A,j_, S_V,j_)). The predicted distribution was obtained by marginalizing over the sensory inputs x_A,j_ and x_V,j_, i.e. internal variables that are not accessible to the experimenter (7). These distributions were generated by simulating x_A,j_ and x_V,j_ in 5000 trials (i.e. continuous variables sampled from Gaussian distributions) for each experimental conditions and inferring spatial estimates S^$^_V,j_ and intensity estimates I^&^_V,j_ from equations (1)-(9) (n.b.: T^(^_V,j_ is not relevant because we did not obtain a response on the timing of the flashes). To link the predicted distributions of participants’ reported number of flashes for flash-beep events one, two and three, we assumed that an observer reported a flash if I^&^_V,j_ was above the perceptual intensity threshold Φ_I_. To link the predicted distributions of the spatial estimates S^$^_V,j_ of perceived flashes to participants’ visual location judgments (i.e., five possible locations), we assumed that participants selected the button that is closest to S^$^_V,j_ and binned S^$^_V,j_ accordingly into a five-bin histogram. If observers did not perceive a flash because the intensity estimate I^&^_V,j_ was below threshold (e.g., the second flash in the invisible rabbit illusion), we counted these simulated trials into a sixth bin of the histogram. We computed these predicted six-bin multinomial distributions for each of the three flash-beep events of a condition, and three distributions separately for each of the experimental conditions. Using the predicted multinomial distributions, we computed the log likelihood of participants’ three flash reports in each condition and summed the log likelihoods across all three flash reports and all experimental conditions of this task.

To obtain maximum likelihood estimates for the 11 parameters of the BCI model (p_common_, σ_S,P_, σ_T,P_, σ_I,P_, σ_s_A, σ_t_A, σ_i_A, σ_s_V, σ_t_V, σ_i_V, Φ_I_), we used a Bayesian optimization algorithm as implemented in the BADS toolbox (43) and initialized this optimization algorithm with multiple different random parameters to prevent local minima. To keep the model simple, spatial, temporal and intensity cues were treated as statistically independent and assumed to carry no specific information in their prior distributions. Consequently, the standard deviations for the spatial, temporal and intensity priors (σ_S,P_, σ_T,P_, σ_I,P_ respectively) were set to large values to mimic uniform distributions (i.e., non-informative priors). To improve the parameter estimation of our four candidate models, we included unisensory condition into an initial parameter estimation: First, we fitted 3 visual parameters (σ_s_V, σ_t_V, σ_i_V) data from six unisensory visual conditions with 50 randomized parameter initialisations, with fixing intensity threshold parameter to 0.5. Plausible lower and upper boundaries were chosen for the fits (Supplementary Table 1), taking the experimental paradigm into account. From this unisensory fit, two parameters (σ_s_V, σ_i_V), related to the cues of the visual stimuli, were fixed. The temporal visual parameter σ_t_V was not fixed, but it was fitted in the multisensory conditions. Because there was no asynchrony in the unisensory visual conditions, fixing σ_t_V based on unisensory visual conditions would severely underestimate the asynchrony effects in visual perception. Finally, since the participants were informed to ignore the sound for the task, and almost perfectly distinguished 2 and 3 beeps in the unisensory auditory precision block, the auditory intensity standard deviation σ_i,A_ was fixed to 10^-10^, reflecting that the participants had very low uncertainty about auditory intensity and nearly perfectly detected the presence or absence of beeps (Supplementary Table 1). The remaining four parameters were then fitted to the AV conditions with 100 random parameter set initialisations, with each optimization iteration simulating 5000 trials with sensory inputs x_A,j_ and x_V,j_. We report the results (i.e., model comparisons and parameters) for models with the highest log likelihood across these initializations (*Table 1 and Supplementary Table 1*). To validate that the models’ parameters can be robustly and accurately estimated from the data using our fitting procedures, we performed a parameter recovery by fitting the model simulated behaviour again. Parameter recovery showed good reproduction of the parameters in our fitting procedure (Supplementary Figure 1).

To identify the optimal model that explains participants’ data, we compared the four candidate models using the Bayesian Information Criterion (BIC; BIC = −2 × LL + m × ln(n), LL = log likelihood, *m* = number of parameters, *n* = number of data points) and Akaike Information Criterion (AIC; AIC = 2 × m − 2 × LL) as an approximation to the model evidence (44). BIC and AIC were aggregated at the group level, i.e. participant-specific BICs and AICs summed over all participants (Table 1). We performed Bayesian model comparison (45) at the random-effects group level as implemented in SPM12 (46) to obtain the protected exceedance probability, which quantifies the probability that a given model is more likely than any other model, beyond differences due to chance (45) for each of the four candidate models. To generate predictions for participants’ flash reports based on the four candidate models, we simulated new x_A,j_ and x_V,j_ for 5000 trials for each experimental condition using the fitted model parameters of each participant. For each simulated trial, we sampled the candidate models’ response from the multinomial predicted distributions and analyzed the models’ responses exactly like participants’ behavioral responses. To reduce the influence random sampling on the models’ predictions, we simulated the tenfold number of trials for each model in each participant.

## ACKNOWLEDGEMENTS

This research was supported by grants from the Deutsche Forschungsgemeinschaft (DFG) to TR (RO 5587/5-1) and DS (SE1859/10-1).

## 8. SUPPLEMENTARY MATERIAL

**Supplementary Table 1.** Fitted parameters for the BCI, FF and FS models. Average parameter values (+-SEM) for the fitted 4 parameters (p_common_, σ_sA_, σ_tA_ and σ_tV_) across 28 participants, as well as the fixed parameters (σ_S,P_, σ_T,P_, σ_I,P,_ σ_sV_, σ_iA_ and σ_tV_) as well as the plausible upper and lower range boundaries for the BADS optimization for the parameter fitting.

| MODELS | $P_{\text{common}}$ | $\sigma_{S,P}$ | $\sigma_{T,P}$ | $\sigma_{I,P}$ | $\sigma_{sA}$ | $\sigma_{tA}$ | $\sigma_{iA}$ | $\sigma_{sv}$ | $\sigma_{tv}$ | $\sigma_{iv}$ |
| --- | --- | --- | --- | --- | --- | --- | --- | --- | --- | --- |
| BCI | 0.76 (± 0.05) | 11.36 | 20 | 1 | 3.41 (± 0.34) | 59.4 (± 5.9) | $10^{-10}$ | 0.84 (± 0.05) | 65.2 (± 4.3) | 0.33 (± 0.01) |
| FF | 1 | 11.36 | 20 | 1 | 3.93 (± 0.31) | 8.2 (± 2.5) | $10^{-10}$ | 0.84 (± 0.05) | 6.7 (± 1) | 0.33 (± 0.01) |
| FS | 0 | 11.36 | 20 | 1 | 0.72 (± 0.29) | 5.2 (± 2.2) | $10^{-10}$ | 0.84 (± 0.05) | 17.1 (± 5.6) | 0.33 (± 0.01) |
| BCI-NP | 0.65 (± 0.05) | 11.36 | 20 | 1 | 3.29 (± 0.37) | 59.5 (± 5.6) | $10^{-10}$ | 0.84 (± 0.05) | 68 (± 4.1) | 0.33 (± 0.01) |
| Boundaries | 0-1 |  |  |  | 0-5.84 | 0-84 |  | 0-1.42 | 0-84 | 0-0.5 |

**Supplementary Figure 1.**
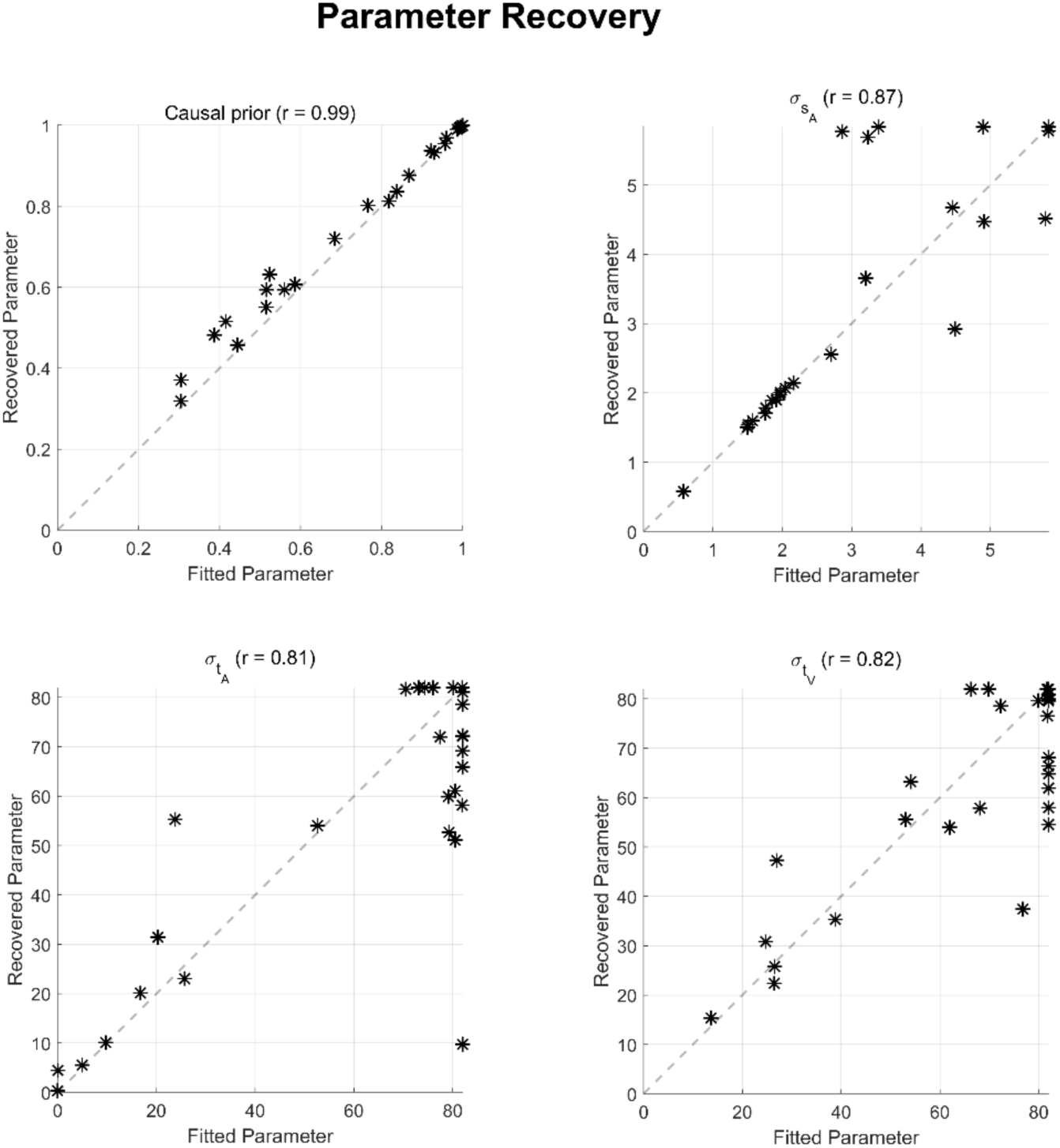
**Parameter recovery for the fitted BCI model parameters (**p_common_, σ_sA_, σ_tA_ and σ_tV_) Recovered parameter values vs. fitted parameter values for the parameters of the BCI model with the correlation coefficients. The black dashed line represents the unit line.

